# Can hyper/hypo-osmoregulating fiddler crabs from the Atlantic coast of South America mobilize intracellular free amino acids as osmotic effectors during salinity challenge?

**DOI:** 10.1101/2022.08.26.505473

**Authors:** Samuel Coelho Faria, John Campbell McNamara

## Abstract

Weakly osmoregulating crustaceans use intracellular free amino acids (FAA) to attenuate cell volume changes consequent to alterations in hemolymph osmolality. Whether semi-terrestrial, strong hyper/hypo-osmoregulators exhibit this ability is not known. We investigate FAA mobilization in muscle tissue of ten fiddler crabs from the genera *Minuca, Leptuca* and *Uca* distributed along the Atlantic coast of South America. Crabs were subjected to severe hypo- or hyper-osmotic challenge at their upper or lower critical salinity limits for five days; control crabs were held in isosmotic media. Hemolymph osmolality was measured, chela muscle FAA were identified and quantified, and percent contribution to intracellular osmolality (%FAA) was calculated. At isosmoticity, total FAA were nominally 2-fold higher in *Minuca* species (≈116 mmol/kg wet mass) than in *Uca* (≈60 mmol/kg wet mass). Glycine, alanine, arginine and taurine constituted >80% of total FAA. On hyper-osmotic challenge, hemolymph osmolalities ranged from 843 to 1,282 mOsm/kg H_2_O. FAA increased, although %FAA remained unaltered. Hypo-osmoregulating crabs thus can mobilize FAA, likely owing to a lesser ability to secrete salt near their upper critical limits. On hypo-osmotic challenge, osmolalities were more tightly regulated, between 475 and 736 mOsm/kg H_2_O. Total FAA and %FAA showed little change, probably due to the crabs’ strong hyper-osmotic extracellular regulation, FAA consequently playing a diminished role in isosmotic intracellular regulation. Total FAA responses to hyper/hypo-osmotic challenge are thus asymmetrical. There was no effect of crab genus on total FAA or on %FAA at isosmoticity or on either osmotic challenge, reinforced by the absence of phylogenetic signal.

## Introduction

Osmoregulation in crustaceans is effected at two distinct levels of structural organization: the systemic level, known as anisosmotic extracellular regulation (AER), and the cellular level, termed isosmotic intracellular regulation (IIR) (Gilles & Péqueux, 1981). The former relies on mechanisms of ion uptake or secretion that maintain hemolymph osmolality within species-specific limits, and is brought about by active ion transport across gill and/or branchiostegite epithelia (Mantel & Farmer, 1983; Péqueux, 1995; Charmantier et al., 2009), antennal glands (Freire et al., 2008) and intestine (McNamara et al., 2005). Such epithelia include structurally specialized ionocytes that possess an elevated mitochondrial volume and exhibit highly amplified membrane surface areas that house a suite of asymmetrically distributed ion-transporting proteins (McNamara et al., 2015). The sodium-proton exchanger, the chloride-bicarbonate exchanger, the sodium-potassium-two chloride symporter and the vacuolar proton ATPase, together with sodium and potassium channels, among others, are located in the numerous evaginations of the apical membrane (Henry et al., 2012). Extensive, deep and narrow invaginations of the basal membrane, intimately associated with mitochondria and continuous with the hemolymph, harbor the sodium-potassium ATPase (Towle & Kays, 1986; Taylor and Taylor, 1992; McNamara & Torres, 1999; Furriel et al., 2010), the sodium-potassium-two chloride symporter and chloride channels, among others (McNamara & Faria, 2012).

Depending on the prevailing osmotic and ionic gradients between an organism’s external medium and its hemolymph, these apical and basal transporters function in a coordinated and complementary manner to bring about net salt uptake or salt secretion across the epithelia, resulting in hyper- or hypo-osmotic regulation, respectively, and constitute short-term, systemic responses. The genes encoding for these transporters are sensitive to external salinity and their expression is up- and/or down-regulated depending on the predominant osmotic gradient (Havird et al., 2013; 2014). To illustrate, often, in marine and estuarine decapods, the genes encoding the α-subunit of the sodium-potassium ATPase and the B-subunit of the vacuolar proton ATPase are up-regulated in low salinities or in fresh water and down-regulated at higher salinities (Mantovani & McNamara, 2021). In freshwater decapods, these genes (Faleiros et al., 2017, 2010) and their proteins (Firmino et al., 2011) are usually down-regulated as salinity increases. Some genes, such as the sodium-potassium-two chloride symporter may be up-regulated in dilute and elevated salinities, revealing a role in both ion uptake and secretion (Maraschi et al., 2021).

Isosmotic intracellular regulation embraces a suite of cellular processes that maintain the intracellular fluid isosmotic with the extracellular fluid, minimizing the cell swelling or shrinkage that accompanies acute volume changes (Claybrooke 1983; (Péqueux, 1995). The active transport of two osmotic components is involved, *i*.*e*., inorganic ions like sodium, potassium, calcium and chloride, and organic osmolytes such as non-essential free amino acids (FAA) like glycine, arginine, alanine, proline and taurine, together with peptides and sugars (Siebers et al., 1972; Gilles & Péqueux, 1981; Pierce, 1982). Ion transport is effected by the sodium-potassium ATPase, sodium-potassium-two chloride symporter, sodium-proton (ammonium) exchanger, and sodium, calcium and chloride channels, among others (Foster et al., 2010, Freire et al., 2013).

The osmotically active FAA pool is regulated by changes in the rates of FAA export to and import from the extracellular fluid, reversible modification in the rates of cellular FAA oxidation and synthesis, and in the rates of protein degradation and synthesis and FAA deamination (McNamara et al., 2004). These two strategies create osmotic gradients down which water may move, usually via the regulated expression of water channels known as aquaporins located in the plasma membrane (Foguesatto et al., 2017; 2019). Isosmotic intracellular regulation perhaps may be a more basal ability, appearing earlier in evolutionary time, likely in prokaryotes or early eukaryotes. The later advent of anisosmotic extracellular regulation would require multicellularity and epithelial specialization (McNamara & Freire, 2022). Together then, mechanisms of AER and IIR preserve those cytosolic conditions optimal for cellular metabolic reactions such as ion, water, enzyme and substrate concentrations.

In species that exhibit modest osmoregulatory capabilities, like many intertidal, marine Brachyura such as *Callinectes danae* (Garçon et al., 2021), intracellular FAAs play a significant role in compensating for large changes in hemolymph osmolality (McNamara, 2022). At low salinity, when hemolymph osmolality becomes reduced, tissue FAA concentrations decrease, diminishing the intra/extracellular osmotic gradient and consequent water influx and cell swelling. At high salinity, tissue FAA increase, reducing cell water loss and cell shrinkage. Diadromous palaemonid shrimps (Augusto et al., 2007a) and freshwater crabs (Augusto et al., 2007b) and squat lobsters (Faria et al., 2011) also show compensatory increases in muscle tissue FAA on salinity challenge, reducing cell water loss to an increasingly osmotically concentrated hemolymph, diminishing cell shrinkage. In contrast, in some strongly hyper/hypo-osmoregulating, intertidal caridean shrimps like *Palaemon northropi*, tissue FAA decrease in low salinity but respond asymmetrically and do not increase in salinities above ambient (Augusto et al., 2009).

In hyper/hypo-regulating semi-terrestrial crabs, like fiddler crabs for example, no information is available on tissue FAA composition or on their putative complementary role in IIR. Fiddler crabs osmoregulate over extensive salinity ranges and can be found in extreme salinity regimes from riverine habitats to hyper-saline salt marshes (Faria et al., 2017; Thurman et al., 2017; Capparelli et al., 2021), often living near their respective lower and upper critical salinities (Capparelli et al., 2022). This raises the question: would FAA and IIR play a role in these crabs that so strongly regulate their hemolymph osmolality by AER and in which tissues would be little challenged osmotically? Might FAA mobilization be asymmetrical on hypo/hyper-osmotic challenge? Further, might such responses be phylogenetically structured overall?

To answer these questions, and given that the role of FAA in IIR in hyper- and particularly in strongly hypo-osmoregulating fiddler crabs is virtually unknown, here we examine the composition and contribution of muscle tissue FAA during severe osmotic challenge in all ten species that occupy osmotic niches ranging from near fresh water to hyper-saline habitats along the Atlantic coast of southeastern South America (Thurman et al., 2013). These findings are contemplated within a phylophysiological context (McNamara and Faria, 2012; Faria et al., 2017), considering current fiddler crab phylogeny (Shih et al., 2016).

## Material and Methods

### Crab collections and laboratory maintenance

We collected ≈70 adult specimens of each of the ten fiddler crab species known to occur in Brazil (Thurman et al., 2013), either directly from the substrate surface by hand or by digging carefully into their burrows with a trowel. The crabs were caught at random during low tide at each locality sampled between June, 2009 (southern winter) and November (early southern summer), 2009.

The fiddler crab species were distributed along the Atlantic coast of South America between the states of Amapá (2° 35’ 47.50” N, 50° 50’ 53.99” W) and Santa Catarina (27° 26’ 57.44” S, 48° 31’ 34.82” W) in Brazil. They included species from three genera: *Minuca burgersi* Holthuis, 1967, *M. mordax* (Smith, 1870), *M. rapax* (Smith, 1870), *M. thayeri* Rathbun, 1900, *M. victoriana* Von Hagen, 1987 and *M. vocator* (Herbst, 1804); *Leptuca cumulanta* Crane, 1943, *L. leptodactyla* Rathbun, 1898 and *L. uruguayensis* Nobili, 1901; and *Uca maracoani* (Latreille, 1802-1803). The range of ≈25° in habitat latitude embraces an extensive variety of ambient conditions (see Thurman et al., 2013 and Faria et al., 2017; 2020 for detailed habitat characteristics), including populations from niches ranging from fresh (*<*0.5 ‰S, 18 mOsm/kg H_2_O) to euhaline seawater (40 ‰S, 1,203 mOsm/kg H_2_O; 1 ‰S = 30 mOsm/kg H_2_O), and found at substrate surface temperatures of from 22 to 36 °C. Surface sediments included coarse to fine grained sand to mud.

The crabs were transported to the Centro de Biologia Marinha, Universidade de São Paulo, in São Sebastião, São Paulo, in closed plastic containers holding sponge cubes moistened with water from each collecting site. In the laboratory, from 20 to 50 crabs of each species were held separately in large plastic boxes (60 cm length × 40 cm width × 30 cm height) at a constant temperature of 25 °C under a natural 12 h light: 12 h dark photoperiod for 3 days prior to the experiments to adjust to laboratory conditions. Each box contained an artificial medium of salinity similar to that of the collection site and was slightly inclined to provide free access by the crabs to a dry surface. Salinities ranged from 3.0 to 17.6 ‰S (90 to 529 mOsm/kg H_2_O) for the *Minuca* species, 12.8 to 20.3 ‰S (385 to 609 mOsm/kg H_2_O) for the *Leptuca* species, and 20.2 ‰S (606 mOsm/kg H_2_O) for *Uca maracoani*. The crabs were not fed during the experiments.

Habitat salinities were measured in the closest available water sources to the collection sites like tide pools and streams. The choice of the closest water source follows previous collection protocols (Thurman, 2002, 2005; Thurman et al., 2010) and enables direct comparison since variability in population habitat salinity is included in the mean habitat salinity for each species.

### Anisosmotic extracellular and isosmotic intracellular regulation

Groups of from four to seven, adult, non-ovigerous, intermolt crabs of either sex, of carapace width >8.0 mm, were used. After adjustment to laboratory conditions in media of salinities similar to those at their collecting sites, the crabs were exposed to three different saline media for 5 days. These media were designed to be either: (i) severely hypo-osmotic [LC_50_, lower critical salinities, 0.6-2.9 ‰S (18-88 mOsm/kg H_2_O) for *Minuca*; 1.7-2.2 ‰S (52-65 mOsm/kg H_2_O) for *Leptuca*; and 6.4 ‰S (193 mOsm/kg H_2_O) for *Uca*]; (ii) approximately isosmotic [19.3-25.5 ‰S (579-765 mOsm/kg H_2_O) for *Minuca*; 24.9-30.4 ‰S (748-912 mOsm/kg H_2_O) for *Leptuca* and *Uca*]; or (iii) severely hyper-osmotic [UC_50_, upper critical salinities, 48.4-82.5 ‰S (1,453-2,475 mOsm/kg H_2_O) for *Minuca*; and 59.5-86.2 ‰S (1,786-2,585 mOsm/kg H_2_O)] for *Leptuca* and *Uca*, depending on the species used (see Table 1 in Faria et al., 2017 for details).

**Table 1.**
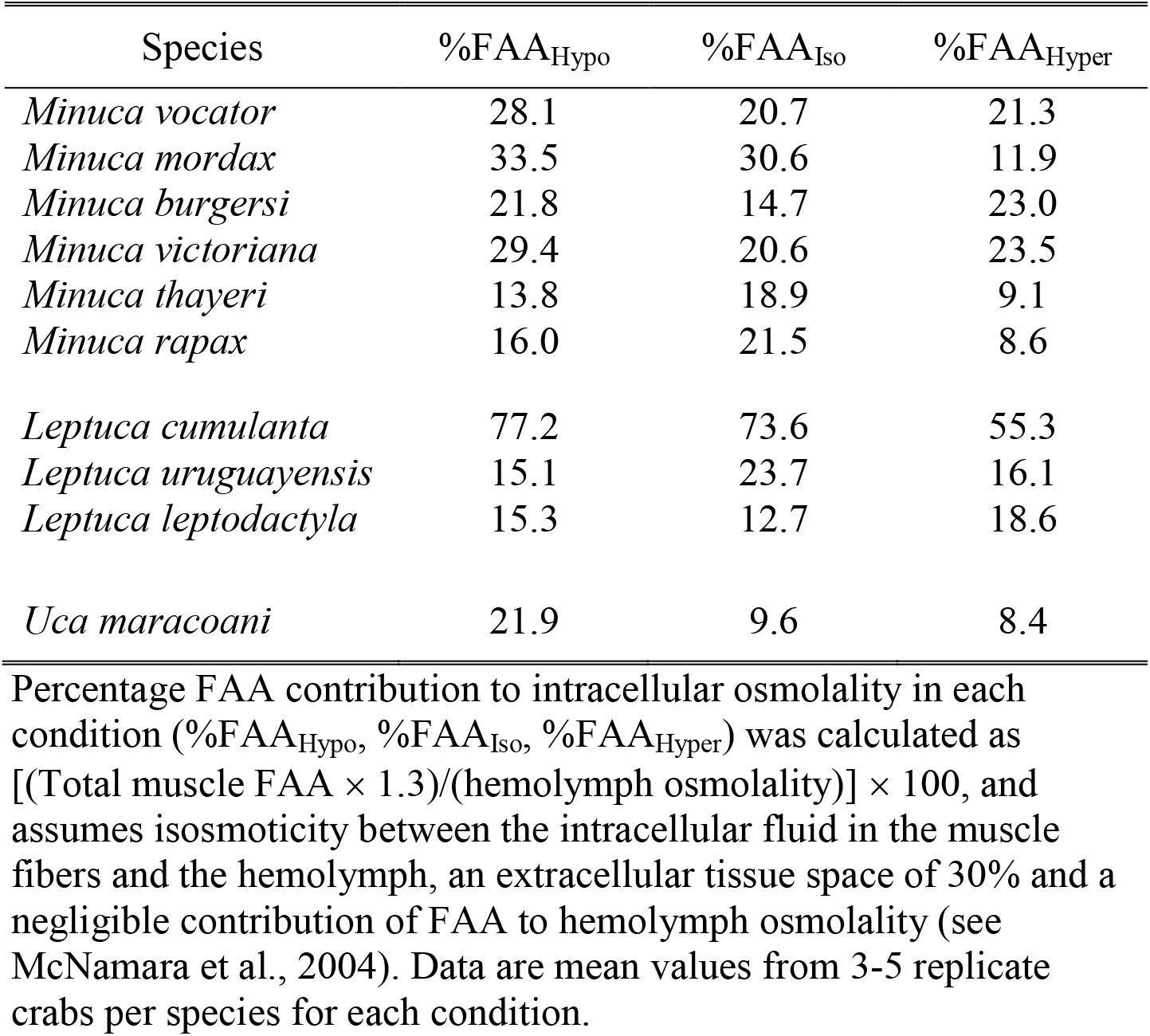
Mean percentage contribution of total free amino acids (%FAA) to muscle tissue osmolality in ten species of fiddler crabs (*Minuca, Leptuca* and *Uca*) from the Atlantic coast of South America. Crabs were held with free access to saline media corresponding to their collection site salinities for 3 days (3.0 to 17.6 ‰S [150 to 529 mOsm/kg H_*2*_O] for the *Minuca* species, 12.8 to 20.3 ‰S [385 to 609 mOsm/kg H_*2*_O] for the *Leptuca* species, and 20.2 ‰S [606 mOsm/kg H_*2*_O] for *Uca maracoani*) to adjust to laboratory conditions. They were then subjected to a severe species-specific hyper- or hypo-osmotic challenge for 5 days. The order of species follows increasing habitat osmolality.

All media were prepared by dissolving artificial sea salts (Instant Ocean, Blacksburg, VA, USA) in distilled water. pH was adjusted to 8.0 using 0.5 to 3 M HCl or NaOH as required (DM-PV pHmeter, Digimed, São Paulo, Brazil).

Crabs were kept under a natural 12 h light: 12 h dark cycle at 25 °C in small covered plastic bowls containing ≈50 mL of each medium. Each bowl held a layer of water of 3-4 mm deep that kept the crabs emerged while permitting voluntary, periodic gill wetting, simulating natural access to water, and allowing evaluation of osmoregulatory ability rather than desiccation tolerance. Mortality was recorded every 24 h when the media were renewed. A crab was considered ‘dead’ if it did not right itself after being turned upside down.

At the end of the 5-day exposure period, a 10-μL hemolymph sample was drawn from the arthrodial membrane between the basis and coxa of the third pereiopod of live crabs using a #25-8 gauge needle coupled to a chilled 1.0-mL syringe. The osmolality of the experimental media and of the hemolymph samples was measured in undiluted, 10-μL aliquots using a vapor pressure osmometer (Wescor 5520 XR, Logan, UT, USA).

Muscle free amino acid (FAA) concentrations were estimated using ≈50 mg muscle tissue dissected from the large chela of each male crab after the 5-day exposure period. The fresh samples were blotted dry uniformly with absorbent paper and their masses measured immediately using a precision balance (±10 μg precision, Ohaus APD 250, Parsippany, NJ, USA). After drying for 24 h in an oven at 50 °C, the samples were reweighed and the initial degree of muscle tissue hydration (% H_2_O) was calculated [(wet mass - dry mass)/wet mass] × 100.

Each sample was then transferred into an individual 0.5-mL Eppendorf tube containing 200 μL distilled water, and homogenized for 1.5 min at 1,500 rpm in crushed ice using a Teflon pestle (Eurostar Power-B, IKA-Werke GmbH & Co., KG, Staufen, Germany). The homogenized samples were centrifuged at 14,000 rpm for 30 min at 25 °C (Fanem Excelsa Baby II, model 206R, São Paulo, Brazil) and the supernatant containing the FAA was quickly transferred to a new Eppendorf tube.

A 20-μL aliquot of supernatant was vacuum dried for 30 min and resuspended in 0.2 M lithium citrate buffer (pH 2.2) containing N-leucine as an internal standard. The FAA analyses were performed in 40-80 μL aliquots of each sample by ion-exchange chromatography using post-chromatographic derivation with ninhydrin (Spackman et al., 1963), employing an amino acid analyzer (Biochrom model 20+, Cambridge, UK).

FAA’s were identified by the retention times of each residue compared to a standard solution (Pierce Amino Acid Standard H, Thermo Fischer Scientific, Waltham, MA, USA) containing 10 nmoles of each amino acid, analyzed under the same conditions. The area of the peak corresponding to each amino acid in the standard was used to calculate the respective concentrations in the samples. The data obtained in nanomoles FAA in 20 μL (per tube) were converted into millimoles FAA per kilogram wet muscle mass using the formula: mmol FAA/kg wet mass = (nmol FAA/20 µL × 10)/[(nmol FAA/mg dry mass) × (100 - %H_2_O)/100].

The percentage contribution of total FAA to intracellular osmolality was estimated using the formula: %FAA contribution = [(Total FAA × 1.3)/(hemolymph osmolality)] × 100, assuming isosmoticity between the intracellular fluid and the hemolymph, an extracellular space of 30% and a negligible contribution of FAA to hemolymph osmolality (McNamara et al., 2004).

### Data fitting, presentation and statistical analyses

Data for hemolymph osmolalities were fitted to quadratic equations using the curve fitting function of SlideWrite Plus for Windows 7 software (Advanced Graphics Software, Inc., Encinitas, CA, USA).

For descriptive and comparative purposes, crab species were ordered between and within their genera according to their decreasing mean habitat salinity, *i*.*e*., *U. maracoani* 20.3 ‰S (606 mOsm/kg H_2_O); *L. leptodactyla* 20.3 ‰S (609 mOsm/kg H_2_O); *L. uruguayensis*, 16.7 ‰S (500 mOsm/kg H_2_O); *L. cumulanta* 12.8 ‰S (385 mOsm/kg H_2_O); *M. thayeri* 17.6 ‰S (539 mOsm/kg H_2_O); *M. rapax* 15.9 ‰S (436 mOsm/kg H_2_O); *M. burgersi* 11.6 ‰S (349 mOsm/kg H_2_O); *M. victoriana* 11.3 ‰S (338 mOsm/kg H_2_O); *M. vocator* 10.2 ‰S (305 mOsm/kg H_2_O), and *M. mordax* 5.0 ‰S (150 mOsm/kg H_2_O) (see Table 1 in Faria et al., 2017).

After testing the data sets for normality of distribution (Kolmogorov-Smirnov test) and equality variance (Levene’s test), the effect of exposure to iso-, hipo- or hiper-osmotic media on hemolymph osmolality and on muscle total free amino acid concentrations in each species was evaluated using a one-way analysis of variance followed by the Student-Newman-Keuls multiple means procedure to locate significantly different means (SigmaStat 2.03 software package, Systat Software Inc., San Jose, CA, USA). Data are given as the mean ± SEM (N). A minimum significance level of α= 0.05 was used throughout, P-values ≤ 0.05 being considered significantly different.

Phylogenetic comparative analyses were performed for IIR traits alone since hemolymph parameters were evaluated previously (Faria et al. 2017). For phylogenetic signal, we used Blomberg’s et al. (2003) *K* statistic, considering the standard error for each species. *K* indicates the level of trait similarity between closely related species pairs. *K* values <1 signify that closely related species resemble each other less than expected across the phylogenetic tree; values >1 suggest that closely related species are more similar than expected under Brownian motion models of evolution. Putative differences among total FAA concentrations consequent to osmotic challenge were tested using a phylogenetic paired t-test (Lindenfors et al., 2010) to control for phylogenetic non-independence of the data, also including the species’ standard errors. The effect of genus was tested using a phylogenetic generalized least squares (PGLS) model, assuming the standard error of each species (Grafen, 1989; Garland & Ives, 2000; Lavin et al., 2008).

All comparative analyses were performed using the *ape* (Paradis et al., 2004), *nlme* (Pinheiro et al., 2012) or *phytools* (Revell, 2012) packages in the R environment (R Development Core Team, 2019), with the minimum significance level set at P = 0.05.

## Results

### Hemolymph osmolality and muscle free amino acids

The hemolymph of all the fiddler crab species held in isosmotic media for 5 days fell between 541 and 783 mOsm/kg H_2_O for the *Minuca* species (Figure 1A), between 739 and 790 mOsm/kg H_2_O for the *Leptuca* species and was largely isosmotic at 908 mOsm/kg H_2_O for *Uca maracoani* (Figure 1B).

**Figure 1.**
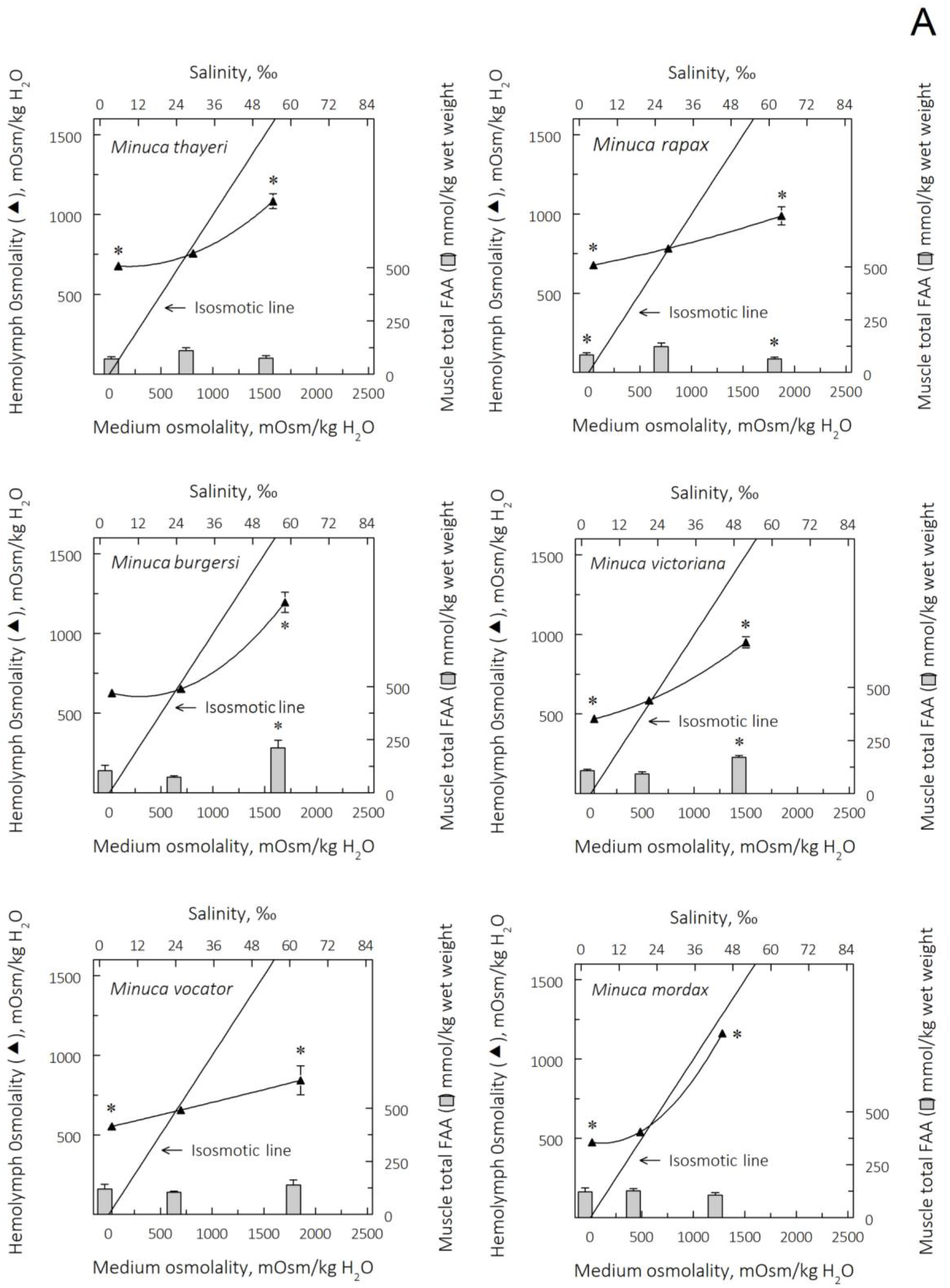
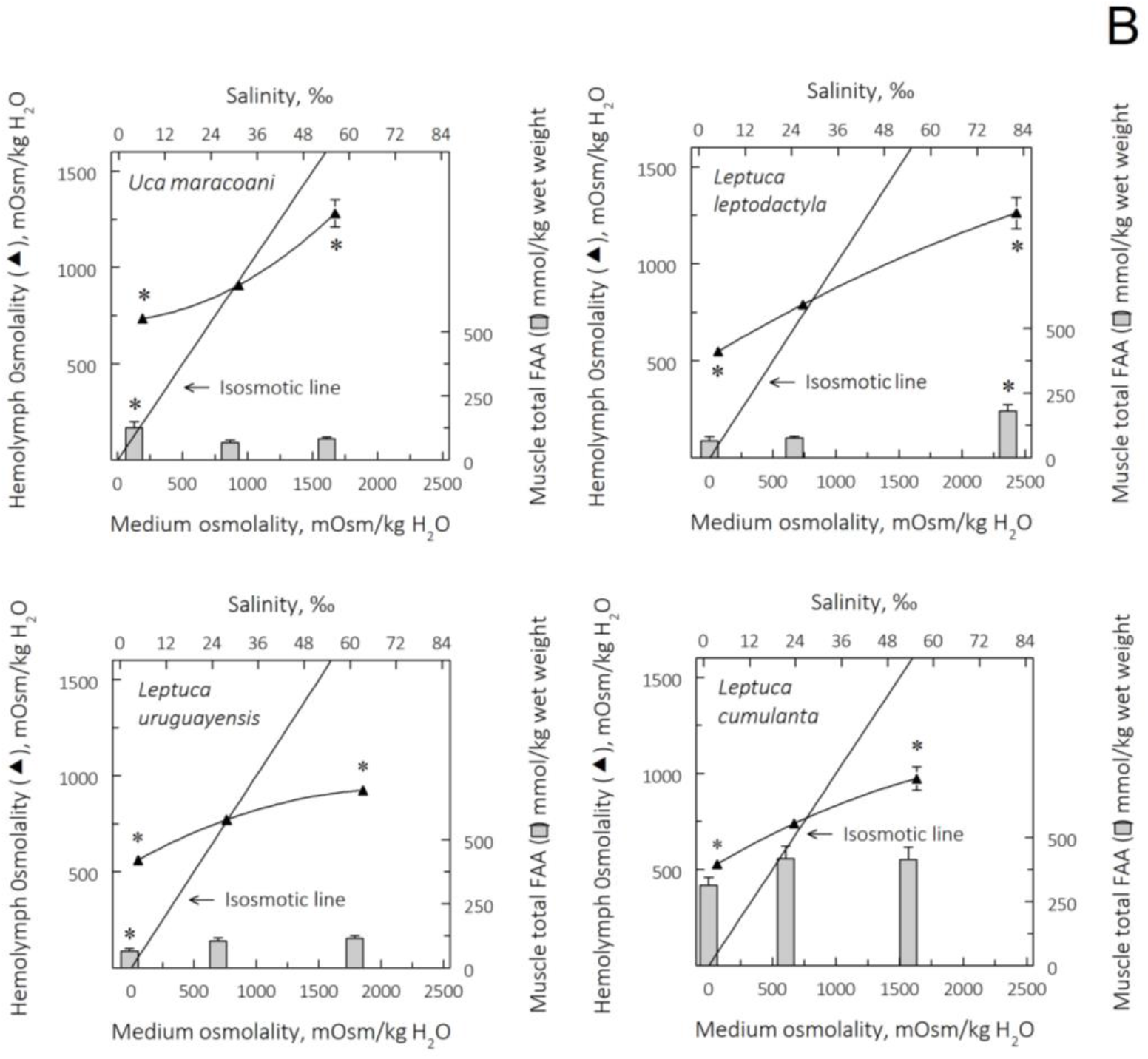
Effect of 5 days acclimation to species-specific isosmotic media (≈25 ‰S, 750 mOsm/kg H_2_O, reference salinity) or species-specific challenge with severely hypo- or hyper-osmotic media on hemolymph osmolality (mOsm/kg H_2_O) and on muscle total free amino acid concentrations (mmol FAA/kg wet mass) in ten fiddler crab species from the Atlantic coast of South America. Prior to osmotic challenge, crabs were held with free access to media of salinity similar to those at their collecting sites for 3 days to adjust to laboratory conditions. Data are the mean ± SEM (N= 4-7) and have been fitted to quadratic equations. Where absent, error bars are smaller than the symbol used. *P≤ 0.05 compared to the isosmotic reference salinity at ≈25 ‰S. **A**, Six species of *Minuca*. **B**, Four species from the genera *Leptuca* and *Uca*.

On hyper-osmotic challenge for 5 days, hemolymph osmolality increased significantly in all species (*Minuca* 843 to 1,195 mOsm/kg H_2_O; *Leptuca* 924 to 1,262 mOsm/kg H_2_O; *U. maracoani* 1,282 ± 70 mOsm/kg H_2_O) (Figures 1A and 1B).

Osmotic differences against the UC_50_ salinities (-Δ) were 117 to 1,011 mOsm/kg H_2_O in the *Minuca* species, 661 to 1,168 mOsm/kg H_2_O in *Leptuca*, and 391 mOsm/kg H_2_O in *U. maracoani*.

After 5-days hypo-osmotic challenge, hemolymph osmolality decreased significantly (ANOVA, SNK, P≤ 0.05) in all *Minuca* species (475 to 678 mOsm/kg H_2_O) except *M. burgersi* (626 ± 8 mOsm/kg H_2_O) (Figure 1A), in all *Leptuca* species (528 to 561 mOsm/kg H_2_O) (Figure 1B) and in *U. maracoani* (736 ± 13 mOsm/kg H_2_O) (Figure 1B). Osmotic differences against the LC_50_ salinities (+Δ) were 436 to 628 mOsm/kg H_2_O in the *Minuca* species, 463 to 509 mOsm/kg H_2_O in *Leptuca*, and 543 mOsm/kg H_2_O in *U. maracoani*.

Muscle total free amino acid (FAA) concentrations in the fiddler crabs after 5 days in isosmotic media were between 74 and 127 mmol/kg wet mass for the *Minuca* species (Figure 1A), between 77 and 418 mmol/kg wet mass for the *Leptuca* species (Figure 1B), and 60 ± 10 mmol/kg wet mass for *U. maracoani* (Figure 1B). After 5-days hyper-osmotic challenge, muscle total FAAs were mainly unaltered, although increasing in *M. burgersi, M. victoriana* (Figures 1A) and *L. leptodactyla* (Figure 1B), but decreasing in *M. rapax* (Figure 1A). Strikingly, FAA concentrations were unaltered in *M. mordax*, despite its virtually isosmotic hemolymph (Figure 1B). After 5-days hypo-osmotic challenge, total FAAs also were mainly unchanged overall, decreasing in *M. rapax* (Figure 1A) and *L. uruguayensis* (Figure 1B), but increasing in *U. maracoani* (Figure 1B) (ANOVA, SNK, P≤0.05).

The mean percentage contribution of total FAAs (%FAA) to muscle tissue osmolality across all species and genera was similar, *i*.*e*., 25% in isosmotic medium, 27% in hypo- and 20% in hyper-osmotic media (Table 1). In isosmotic media, mean %FAA contributions were 21.2 ± 2.1 % in *Minuca*; 36.7 ± 18.7 in *Leptuca*, and 9.6 in *Uca*. In hyper-osmotic media contributions were 16.2 ± 2.9 % in *Minuca*; 30.0 ± 12.7 % in *Leptuca* and 8.4 % in *Uca*. In hypo-osmotic media, 23.8 ± 3.2 % in *Minuca* species; 35.9 ± 20.7 % in *Leptuca*; and 21.9 % in *Uca*.

Compared to isosmoticity, in hyper-osmotic media, mean %FAA increased by ≈30% in *M. vocator, M. burgersi, M. victoriana* and *L. leptodactyla*; in the remaining species, %FAA decreased unexpectedly by ≈40%, and by ≈60% in *M. mordax* (Table 1). In hypo-osmotic media, mean %FAA decreased by ≈30% in *M. thayeri, M. rapax* and *L. uruguayensis*; incongruously, in all other species, mean %FAA increased by ≈40% (Table 1).

Glycine, alanine, arginine and taurine constituted ≈80% of the muscle total FAA pool in all fiddler crab species. Changes in the concentrations of these individual FAAs underpinned the alterations seen in all total FAA concentrations significantly affected by osmotic challenge (Table 2).

**Table 2.**
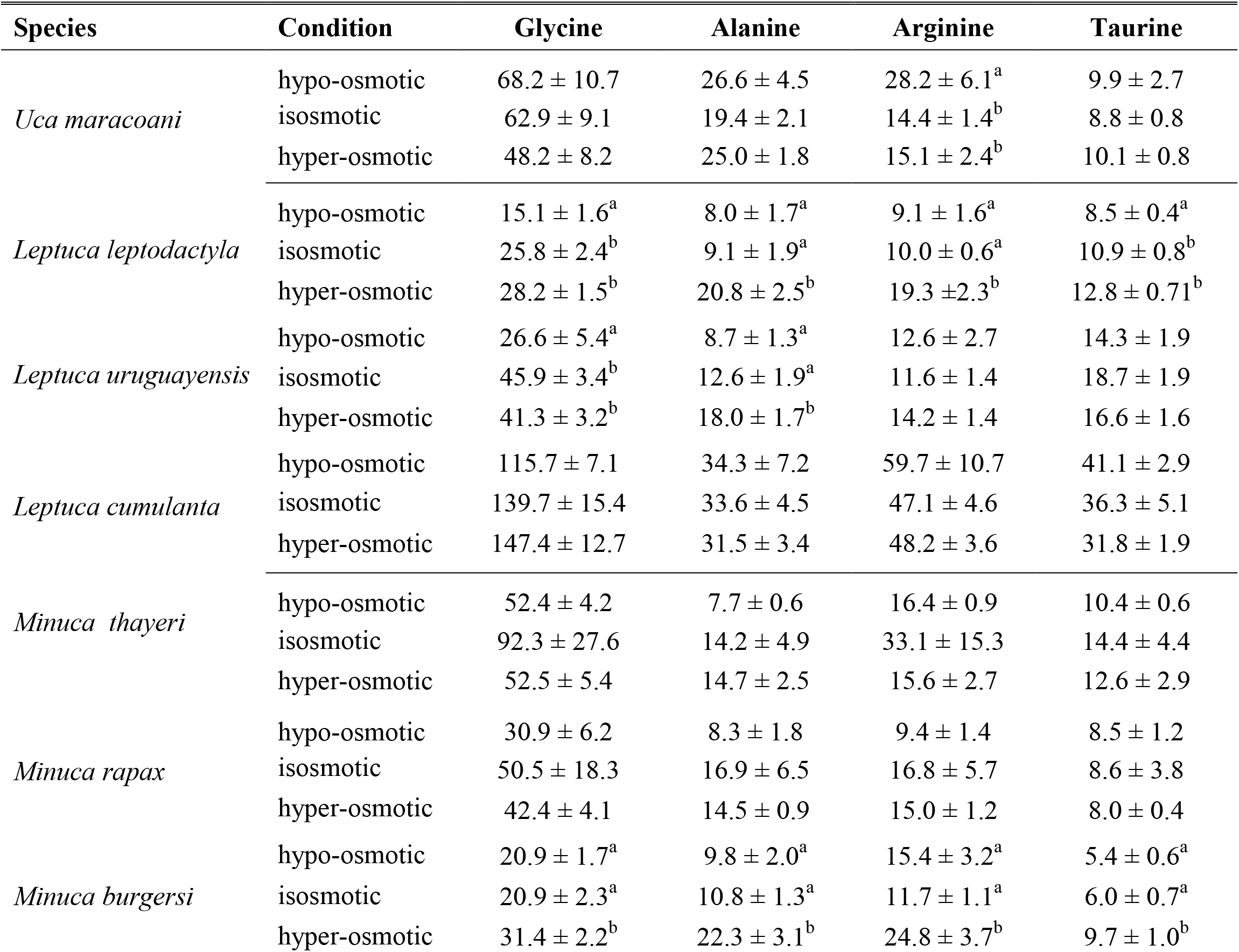
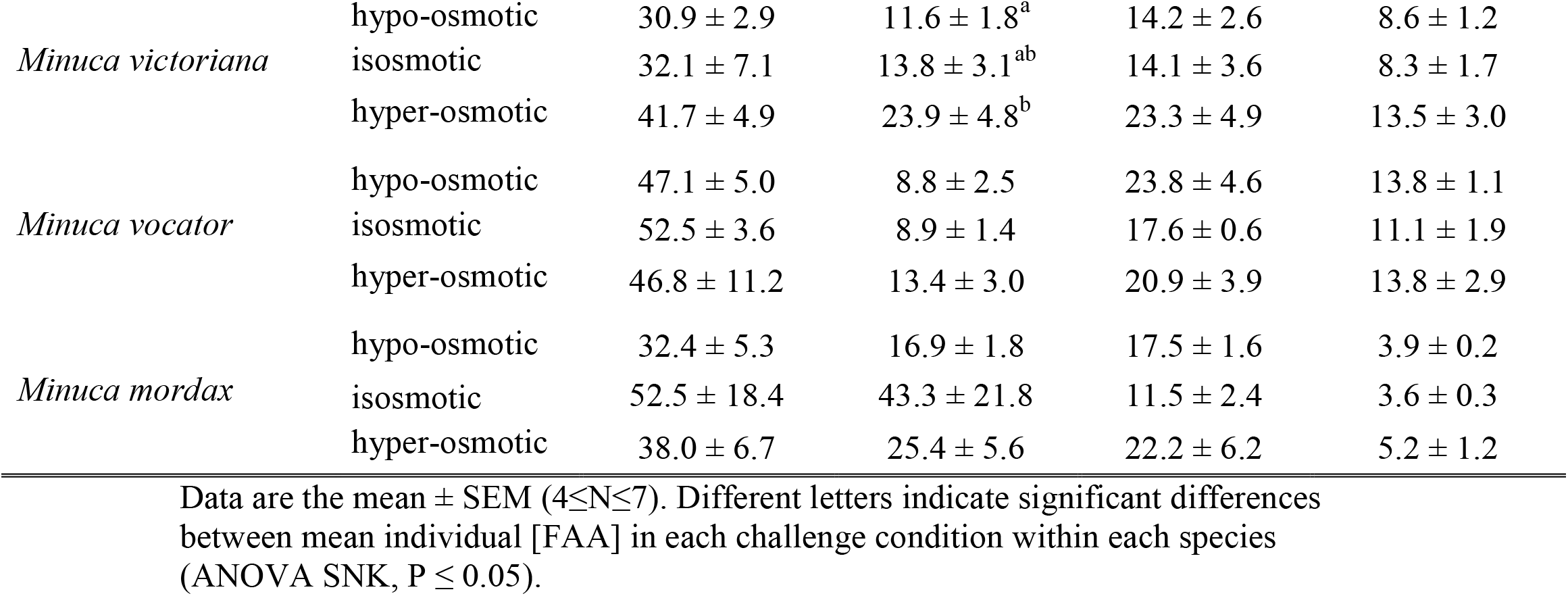
Concentrations of the predominant free amino acids (FAA, mmol FAA/kg wet mass) in the chela muscle of ten species of fiddler crabs (*Uca, Leptuca* and *Minuca*) from the Atlantic coast of South America. Crabs were held with free access to saline media corresponding to their collection site salinities (3.0 to 17.6 ‰S [150 to 529 mOsm/kg H_2_O] for the *Minuca* species, 12.8 to 20.3 ‰S [385 to 609 mOsm/kg H_2_O] for the *Leptuca* species, and 20.2 ‰S [606 mOsm/kg H_2_O] for *Uca maracoani*) for 3 days to adjust to laboratory conditions. They were then subjected to severe species-specific hypo- or hyper-osmotic challenges, or to their respective isosmotic salinities, for 5 days. Glycine, alanine, arginine and taurine constitute more than 80% of the total FAA pool.

### Comparative phylogenetic analyses among genera and species

The absence of a genus effect on total muscle FAA and on %FAA contribution to muscle tissue osmolality at isosmoticity or under hyper- or hypo-osmotic challenge (pGLS, 0.47≤ F ≤0.14; 0.65≤ P≤0.85) is substantiated by the lack of phylogenetic signal (Blomberg’s K, 0.38≤ K ≤0.47; 0.38≤ P ≤0.49). Thus, closely related fiddler crabs species do not share similar total muscle FAA or %FAA.

There were no species related differences among total muscle FAA on hypo-osmotic challenge for 5 days (phylogenetic paired t-test, t=0.76, P=0.47), or among %FAA contributions to muscle tissue osmolality (t=1.47, P=0.19). However, on hyper-osmotic challenge, muscle FAA increased compared to isosmoticity (t=-7.94, P≤0.001), although %FAA were unaltered (t=1.79, P=0.12).

## Discussion

Fiddler crab species from the Atlantic coast of South America survive remarkably well in a widely varying range of saline environments, mainly a consequence of their ability to strongly regulate hemolymph osmolality (Faria et al., 2017; Thurman et al., 2017). Despite some alteration in hemolymph osmotic concentrations, here we show that on hypo-osmotic challenge at their lower critical salinities, all ten fiddler crabs exhibit low hemolymph osmolalities and very strong (27: 1) to moderate (4: 1) hyper-osmotic gradients. On hyper-osmotic challenge at their upper critical salinities, hemolymph osmolalities are elevated and show notable variability, also exhibiting from strong (0.45: 1) to weak (0.91: 1) hypo-osmotic gradients. While mean total muscle FAA increased on hyper-osmotic challenge, %FAA contribution to tissue osmolality remained unaltered. The absence of a genus effect on FAA concentration and on %FAA is reinforced by the lack of significant phylogenetic signal. Thus, while interspecific variability in hemolymph osmolality of South Atlantic fiddler crabs is similar overall, hyper/hypo-osmoregulatory ability is asymmetrical, the crabs exhibiting stronger hyper-regulatory capability, *i*.*e*., the ability to uptake salt in dilute media. Such asymmetry extends to the mobilization of FAA, limited to hyper-osmotic challenge.

No pattern of change in muscle total FAA was discernible on osmotic challenge. Total FAA concentrations were insensitive to osmotic challenge in *L. cumulanta, L. thayeri, M. vocator* and *M. mordax*, increased on hyper-osmotic challenge in *L. leptodactyla, M. burgersi* and *M. victoriana*, and decreased on hypo-osmotic challenge in *L. uruguayensis* and *M. rapax*. Total FAA responses were incongruous in *M. rapax* and *U. maracoani*, increasing in hypo-osmotic media. The oligohaline, biting fiddler crab, *Minuca mordax*, is an illustrative case in point. Despite an unusual 2.2-fold increase in hemolymph osmolality on hyper-osmotic challenge, muscle total FAA concentration was unaltered (Fig. 1A) while %FAA contribution to intracellular osmolality decreased by 60% (Table 1). These findings suggest that total FAA are not mobilized in a predictable manner in hyper/hypo-osmoregulating ocypodids in contrast to weakly hyper-osmoregulating portunids like *Callinectes danae* (McNamara, 2022), hyper-osmoregulating diadromous palaemonids such as *Macrobrachium olfersi* and *M. amazonicum* (Augusto et al., 2007a) or hololimnetic freshwater dwellers like the trichodactylid crab *Dilocarcinus pagei* (Augusto et al., 2007b), aeglid squat lobster *Aegla franca*, and *M. brasiliense* (Faria et al., 2011).

In contrast to the predicted responses, *i*.*e*., that alterations in percentage total FAA contribution to intracellular osmolality might buffer cell volume changes, no clear pattern could be established either. Mean %FAA contributions for all species were 25% at isosmoticity, 20% on hyper-, and 27% on hypo-osmotic challenge, at variance with the notion that relative FAA contributions increase on hyper-osmotic challenge and diminish on hypo-osmotic challenge, buffering respective cell volume decreases and increases. To illustrate, compared to isosmoticity, %FAA contribution to intracellular osmolality *decreases* incongruously by ≈60% on hyper-osmotic challenge in *M. rapax, M. thayeri* and *M. mordax*, and by ≈30% in *L. cumulanta* and *L. uruguayensis*. On hypo-osmotic challenge, %FAA contribution *increases* discordantly by ≈35% in *M. vocator, M. mordax, M. burgersi* and *M. victoriana*, by ≈12% in *L. cumulanta* and *L. uruguayensis*, and by ≈130% in *U. maracoani*. Both sets of responses stand in stark contrast to the predicted effects and neither shows phylogenetic signal.

Although lacking an effect of genus, at isosmoticity, total muscle free amino acids (FAA) were nominally higher in the *Minuca* species (≈116 *cf*. ≈95 mmol/kg wet mass in *Leptuca*), although disparately high in *L. cumulanta* alone (390 mmol/kg wet mass); *Uca maracoani* showed the lowest titers overall (≈60 mmol/kg wet mass), unusual for an infralittoral species. These values lie well below those for various marine Brachyura (253-461 mmol/kg wet mass) including *Leptuca pugilator* (233 mmol/kg wet mass), Astacidea (278-530 mmol/kg wet mass) and Dendrobranchiata (199-320 mmol/kg wet mass) (Faria et al., 2011).

Among the main FAAs that constitute more than 80% of the fiddler crabs’ total muscle FAA pool are glycine, alanine, arginine and taurine. Some individual FAAs are mobilized in a predictable manner in a few species, *e*.*g*., on hyper-osmotic challenge, alanine and arginine in *L. leptodactyla*, all FAA in *M. burgersi*, and alanine in *L. uruguayensis* and *M. victoriana* increase; on hypo-osmotic challenge, glycine and taurine in *L. leptodactyla* and glycine in *L. uruguayensis* decrease. These are minor contributions, however, and no clear pattern prevails. FAA mobilization on hyper-osmotic challenge is thought to result from increased glutamate dehydrogenase activity owing to extracellular ionic effects, favoring amination of α-ketoglutarate to glutamate, a proline precursor and an amino-group donor for glycine and alanine (Bishop & Burton, 1993). Such a mechanism likely would be phylogenetically conserved, *i*.*e*., closely related species should exhibit more similar FAA concentrations and possibly enzyme kinetics than distantly related species.

However, we show here that muscle tissue FAA concentrations and their contribution to cellular osmolality are not phylogenetically structured, and further, that there is no effect of genus, which suggests a role for the crabs’ osmotic niches in having shaped interspecific FAA variability. The nature of this variability is plastic, suggesting relaxed selection pressure on mechanisms of intracellular FAA metabolism. Thus, the lack of a species related pattern in FAA mobilization appears to derive from the crabs’ excellent overall capacity for anisosmotic extracellular regulation. Mean total FAA concentration increased on hyper-osmotic challenge, demonstrating mobilization of intracellular osmotic effectors when hemolymph osmolality increases. However, on hypo-osmotic challenge, intracellular FAA are not reduced owing to lesser decreases in hemolymph osmolality due to strong hyper-regulatory capability.

This scenario diverges from an evolutionary pattern established for crustaceans (McNamara & Faria, 2012) in which total muscle FAA titers correlate positively with hemolymph osmolality (R = +0.82, P <0.0001) across the main decapod sub/infraorders, *i*.*e*., marine taxa exhibit higher FAA concentrations than freshwater representatives. The overall low FAA concentrations found in South American fiddler crabs (≈110 mmol/kg wet mass) are more reminiscent of freshwater taxa like the Caridea, Brachyura and Astacidea.

We conclude that hyper/hypo-osmoregulating fiddler crabs from the Atlantic shores of South America mobilize FAA asymmetrically. Species of *Minuca, Leptuca* and *Uca* employ muscle FAA during hyper-osmotic challenge while on hypo-osmotic challenge, a role for FAA is less evident owing to efficient anisosmotic extracellular regulation that maintains hemolymph osmolality within exacting limits.

## Acknowledgments

This article is dedicated to Professor Lewis Joel Greene, founder and director of the Centro de Química de Proteínas da Fundação Hemocentro de Ribeirão Preto, to whom we are indebted for access to amino acid analysis facilities over many years, including the present investigation. JCM and SCF are also beholden to Professor Greene for his instructive insights into the role of amino acids and his guidance in our better understanding molecular cellular function.

We express our sincere gratitude to Professor Carl Thurman, Department of Biology, University of Northern Iowa, for his comradeship in the field and laboratory, for his enthusiastic teachings on fiddler crab biology, and his accumulated wisdom in collecting, identifying and transporting the species of fiddler crabs used here.

We are most grateful to the Centro de Biologia Marinha, Universidade de São Paulo, for providing laboratory space and facilities (Project #2009/06). We thank Claudia Antunes, Mariana Capparelli, Rogério Faleiros, Melanie Hopkins, Sarah Milograna and Elaine Ribeiro for assistance with field work. Crabs were collected under permits #18559-1, #23976-1, #29594-1 and #29594-3 to JCM from the Instituto Chico Mendes de Conservação da Biodiversidade (IBAMA/MMA).

This investigation constitutes part of an M Sc dissertation submitted by SCF to the Graduate Program in Comparative Biology, Departamento de Biologia, FFCLRP, Universidade de São Paulo.

## Ethical considerations

This investigation complies with all local, state, federal and international guidelines as regards the use of invertebrate animals in scientific research. This study also complies with the ARRIVE guidelines.

---

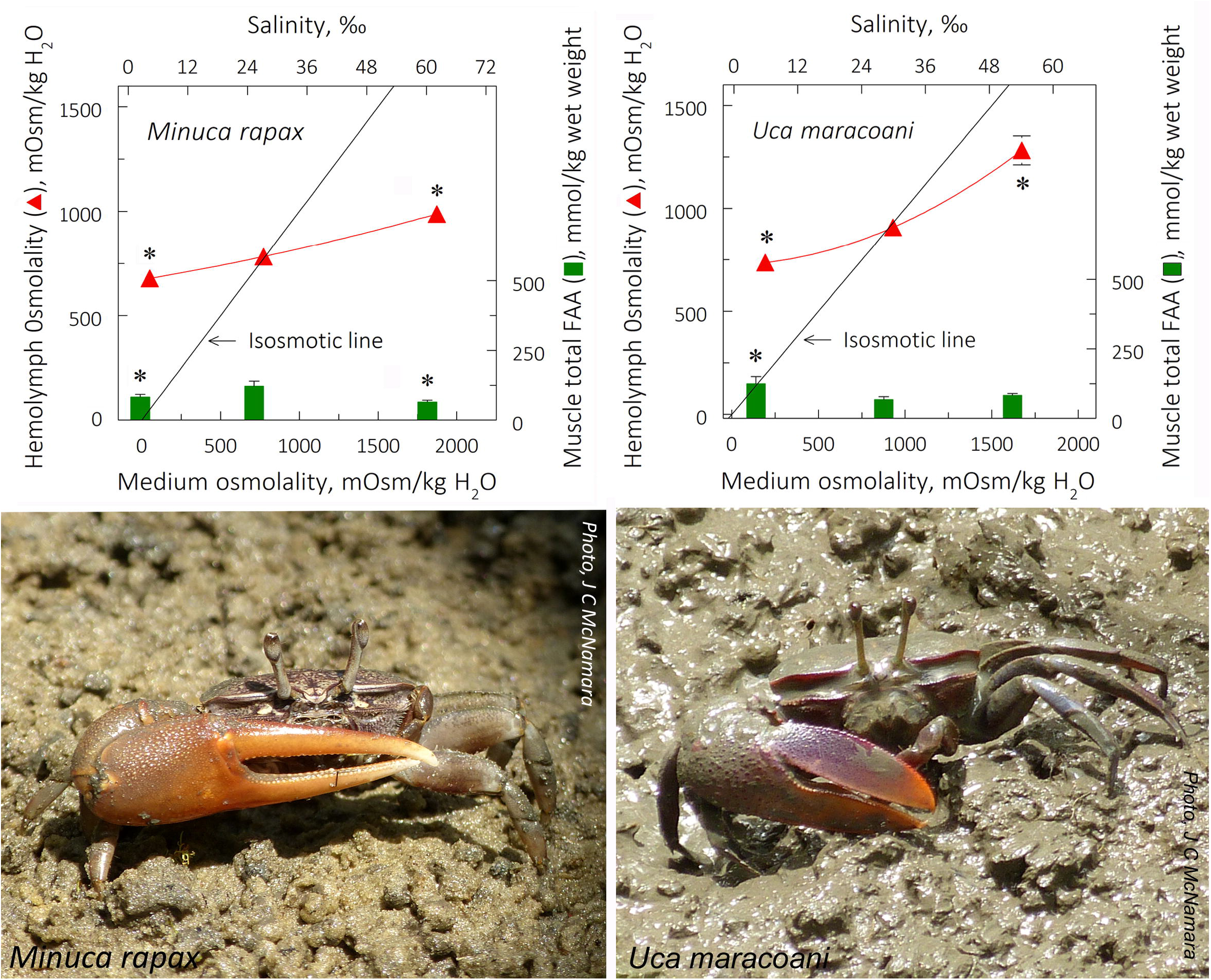

